# Acidic pH-dependent depletion of *Mycobacterium tuberculosis* thiol pools potentiates antibiotics and oxidizing agents

**DOI:** 10.1101/095448

**Authors:** Garry B. Coulson, Benjamin K. Johnson, Christopher J. Colvin, Robert J. Fillinger, Huiqing Zheng, Elizabeth R. Haiderer, Neal D. Hammer, Robert B. Abramovitch

**Affiliations:** Department of Microbiology and Molecular Genetics, Michigan State University, East Lansing, Michigan, 48824, United States.

**Keywords:** *Mycobacterium tuberculosis*, redox homeostasis, oxidative stress, environmental adaptation.

## Abstract

*Mycobacterium tuberculosis* (Mtb) must sense and adapt to immune pressures such as acidic pH and reactive oxygen species (ROS) during pathogenesis. The goal of this study was to isolate compounds that inhibit acidic pH resistance, thus defining virulence pathways that are vulnerable to chemotherapy. Here we report that the acidic pH-dependent compound AC2P36 depletes intracellular thiol pools, sensitizes Mtb to killing by acidic pH, and potentiates the bactericidal activity of isoniazid, clofazimine, and oxidizing agents. We show that the pH-dependent activity of AC2P36 is associated with metabolic stress at acidic pH and a pH-dependent accumulation of intracellular ROS. Mechanism of action studies show that AC2P36 directly depletes Mtb thiol pools. These data support a model where chemical depletion of Mtb thiol pools at acidic pH enhances sensitivity to oxidative damage, resulting in bacterial killing and potentiation of antibiotics.

## Introduction

*Mycobacterium tuberculosis* (Mtb), the etiologic agent of tuberculosis (TB), remains a major challenge in global health. Despite concerted efforts and significant resources dedicated to controlling this disease, an estimated 10.4 million illnesses and 1.8 million deaths occurred worldwide in 2015 (WHO, 2016). A major obstacle in TB control and eradication strategies is the lack of a uniformly effective vaccine and the emergence of drug-resistant TB. Multidrug-resistant TB infections are responsible for approximately 500,000 new cases annually (WHO, 2016). The emergence of drug-resistance significantly undermines global progress in TB control and underscores the importance of developing new drugs with novel targets or mechanisms of action.

Long-term intracellular survival of Mtb in host macrophages is dependent on the organisms’ ability to adapt to and resist numerous host environmental stresses encountered during infection (e.g. oxidative, nitrosative and acid stresses). In resting macrophages, Mtb inhibits phagosome maturation and prevents phagolysosome fusion. Exclusion of the vacuolar proton-ATPase in the Mtb-containing vacuole results in a mildly acidic phagosomal pH (pH 6.1- 6.4) (MacMicking, et al., 2003; Sturgill-Koszycki, et al., 1994). Classical activation of the macrophage by IFN-γ removes this Mtb-mediated blockade allowing the phagosome to fully acidify (pH 4.5 - 5.4) (MacMicking, et al., 2003; Schaible, et al., 1998; Via, et al., 1998). Gene expression profiling has demonstrated that pH-responsive genes are induced in macrophages, signifying that Mtb encounters an acidic environment during *in vitro* macrophage infection (Fisher, et al., 2002; Rohde, et al., 2007; Rohde, et al., 2012). This acidic pH regulon closely resembles the *phoPR* two-component regulatory system (TCS) regulon (Abramovitch, et al., 2011; Gonzalo-Asensio, et al., 2008; Rohde, et al., 2007; Walters, et al., 2006), suggesting that *phoPR* may play a role in pH-driven adaptations. Further evidence that Mtb resides in acidified environments comes from the observation that the TB drug pyrazinamide has enhanced activity against Mtb at acidic pH (Zhang and Mitchison, 2003; Zhang, et al., 2003).

Survival of *M. tuberculosis* at acidic pH during infection requires maintenance of cytoplasmic pH-homeostasis. Mtb can resist phagolysosomal concentrations of acid when grown in liquid broth buffered at pH 4.5-5.0 and is also capable of maintaining its cytoplasmic pH in IFN-γ –activated macrophages, suggesting that Mtb has robust protective mechanisms for acid resistance (Vandal, et al., 2008; Zhang, et al., 1999). One of the Mtb determinants required for acid resistance and maintenance of pH homeostasis is a membrane-associated serine protease (Rv3671c) (Vandal, et al., 2008). Mutants of Rv3671c are unable to maintain neutral cytoplasmic pH either in acidic growth medium or in the phagolysosomes of IFN-γ-activated macrophages (Vandal, et al., 2008). A pore-forming protein known as OmpA is also important for acid resistance (Raynaud, et al., 2002). *ompA* expression is induced at pH 5.5, and mutants in this gene are attenuated for growth in macrophages and mice (Raynaud, et al., 2002). Lastly, a putative magnesium transporter (MgtC) was also found to be essential for growth *in vitro* at a mildly acidic pH of 6.25, but only in the presence of low Mg^2+^ concentrations (Buchmeier, et al., 2000). Disruption of MgtC resulted in attenuated growth in macrophages and mice, suggesting that Mg^2+^ acquisition becomes important when Mtb is exposed to the low pH of the phagosomal compartment (Buchmeier, et al., 2000).

Acidic pH causes widespread changes to Mtb physiology, including the induction of numerous stress genes and the PhoPR regulon. Additionally, when Mtb is grown on single carbon sources at acidic pH, Mtb requires carbon sources that fuel the anaplerotic node, including pyruvate, acetate and oxaloacetate, demonstrating that metabolism is remodeled at acidic pH (Baker, et al., 2014). Acidic pH also leads to alterations in intracellular redox status, resulting in a reducing environment within the Mtb cytoplasm and in the absence of the pH-regulated TCS PhoPR, the cytoplasm is reduced further at acidic pH, indicating that pH-dependent mechanisms exist to maintain redox homeostasis. Indeed, the redox sensor WhiB3 (which co-regulates PhoPR regulated genes such as *pks2)* and the redox buffer mycothiol are required to maintain redox homeostasis at acidic pH and during macrophage infection (Mehta, et al., 2016; Singh, et al., 2009). Therefore, surviving acid stress and redox homeostasis are associated physiologies, although the mechanisms driving pH-dependent reductive stress remain poorly characterized.

Mtb is exposed to acidic pH at various stages during infection and this acidic environment may act as a critical cue for the initiation of various adaptive mechanisms required for survival in the host. Therefore, the goal of this study was to identify and characterize compounds that selectively inhibit Mtb growth at acidic pH. These compounds may identify novel drug targets or pathways that are essential for growth at acidic pH, which would have been missed in previously reported drug screens performed in rich medium at neutral pH. Our team has completed two, independent high throughput screens searching for inhibitors of acidic pH-inducible or hypoxia-inducible signaling pathways (Johnson, et al., 2015; Zheng, et al., 2016). The screens were performed under near identical conditions, including the same ~220,000 compound library, with the primary difference being the pH of the medium (pH 7.0 or pH 5.7). Comparisons of hits between the two screens revealed several compounds that inhibit Mtb growth at acidic pH, but have no impact at neutral pH. We hypothesized that pH-dependent compounds may be targeting pathways that are only essential for growth at acidic pH. Here we report that one of the pH-dependent compounds identified in the screen, named AC2P36, kills Mtb at acidic pH, depletes intracellular thiol pools, promotes the accumulation of intracellular ROS and potentiates the activity of isoniazid, clofazimine and oxidizing agents. These findings support that redox homeostasis pathways are vulnerable pH-dependent targets and when inhibited, promote Mtb killing and potentiation of antimicrobials.

## Results

### Discovery of AC2P36 as an acidic pH-dependent inhibitor of Mtb growth

Two whole-cell phenotypic high throughput screens (HTS) were conducted at neutral and acidic pH for inhibitors of the DosRST and PhoPR regulons, respectively, using an identical chemical library (Johnson, et al., 2015; Zheng, et al., 2016). The ~220,000 compound screening library is composed of small molecules representing broad chemical diversity. By examining the overlap among inhibitors that reduce Mtb growth, compounds were found that selectively inhibited Mtb growth at acidic pH. Small molecules that exhibited a profile of at least 50% growth reduction and were unique to the HTS at acidic pH were considered for additional follow-up and characterization as pH-dependent inhibitors of Mtb growth. AC2P36 (5-chloro-N-(3-chloro-4-methoxyphenyl)-2-methylsulfonylpyrimidine-4-carboxamide, Fig. 1A) was identified as a compound that met the profile of slowing Mtb growth at acidic pH while leaving growth at neutral pH unaffected. At pH 5.7, AC2P36 inhibits Mtb growth with a half-maximal effective concentration (EC_50_) of ~3 μM, whereas at pH 7.0 the EC_50_ is >30 μM (Fig. 1B), thus AC2P36 exhibits ~10-fold selectivity at acidic pH. At concentrations >30 μM, Mtb growth is slowed at neutral pH suggesting that the mechanism through which AC2P36 is acting may not be exclusive to acidic pH. AC2P36 does not modulate Mtb cytoplasmic pH, indicating it does not function as an ionophore (Supplemental Figure 1A). To assess whether AC2P36 acts in a bactericidal or bacteriostatic manner at acidic pH, time-dependent and dose-dependent survival assays were undertaken. AC2P36 at a concentration of 20 μM exhibits time-dependent killing causing a progressive, ~100-fold reduction of viability over the course of 7 days (Fig. 1C). AC2P36 bacteriocidal activity is dose-dependent with bactericidal activity observed at concentrations >10 μM following 6 days of treatment (Fig. 1D). Mtb exhibits a reduced cytoplasm at acidic pH and requires PhoPR to maintain redox poise (Baker, et al., 2014). To test whether the ability of AC2P36 to kill Mtb at acidic pH is the result of altered redox homeostasis, Mtb expressing a redox active GFP was treated with increasing concentrations of AC2P36. AC2P36 treatment leads to a reduced cytoplasm in a dose-dependent and pH-independent manner (Fig. 1E). Given that PhoPR has been shown to be required for pH-dependent maintenance of redox homeostasis we hypothesized that a *∆phoPR* mutant may be more susceptible to AC2P36, however, no differences were observed between WT or *∆phoPR* sensitivity to AC2P36 at acidic pH (Fig. 1F). These data demonstrate that AC2P36 inhibits Mtb growth in a pH-dependent manner, is bactericidal, and promotes a reduced cytoplasm.

**Figure 1.**
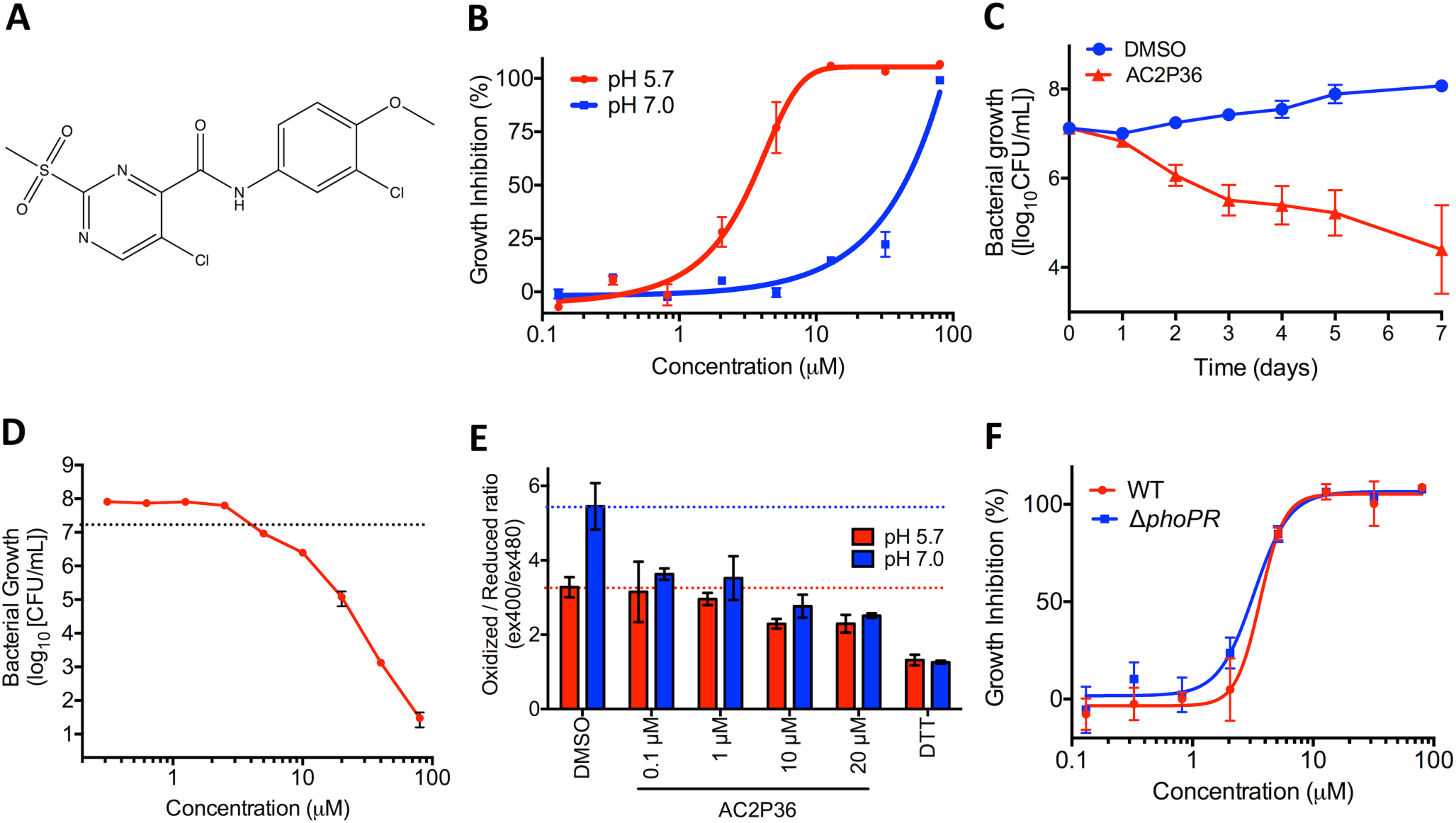
AC2P36 is a pH-selective, bacteriocidal compound that promotes a reduced intracellular environment. A) Chemical structure of AC2P36 (5-chloro-N-(3-chloro-4-methoxyphenyl)-2-methylsulfonylpyrimidine-4-carboxamide). B) Treatment with AC2P36 inhibits Mtb growth in a dose-dependent manner at pH 5.7 with an EC_50_ of 3.2 μM. Bacterial growth is inhibited at pH 7.0 in a dose-dependent manner at concentrations greater than 30 μM. C) Timedependent killing of Mtb at pH 5.7 treated with 20 μM AC2P36. D) Dose-dependent killing of Mtb at pH 5.7 following 6 days of treatment with AC2P36. The dotted line indicates the CFUs of the initial inoculum. E) AC2P36 promotes a reduced cytoplasm in a dose-dependent and pHindependent manner. Cytoplasmic redox status was measured using redox-sensitive ro-GFP. F). The CDC1551 *(AphoPR)* has similar sensitivity to AC2P36 as WT CDC1551. Data shown are representative of at least two independent experiments and error bars represent the standard deviation of the mean.

### AC2P36 exhibits a narrow spectrum of activity

To assess whether AC2P36 acts as a broad inhibitor of bacterial growth at acidic pH, selected Gram-positive and Gram-negative bacteria were treated with the compound at neutral and acidic pH (Supplemental Table 1). Only the Gram-positive species *Staphylococcus aureus* and *Enterococcus faecalis* exhibit susceptibility to AC2P36 (Supplemental Table 1), whereas the Gram-negative bacterial species *Escherichia coli, Pseudomonas auruginosa,* and *Proteus vulgaris* were relatively unaffected. Notably, in both *S. aureus and E. faecalis,* AC2P36 sensitivity was pH-independent and the compound was less potent. Together, these data suggest Mtb is selectively sensitive to AC2P36, but that a common pathway may be targeted in Mtb, *S. aureus* and *E. faecalis*.

### AC2P36 causes a thiol-associated stress response

To further characterize the mechanism of action of AC2P36, global transcriptional profiling was performed to identify pathways that are differentially expressed in response to AC2P36 treatment. Mtb was inoculated into rich medium buffered at pH 7.0 or 5.7 in the presence of either 10 μM AC2P36 or an equivalent volume of DMSO. After 4 hours incubation, total RNA was isolated and subjected to RNA sequencing. AC2P36 treatment of Mtb caused the induction of 167 genes (>2-fold, q<0.05) and 180 genes (>2-fold, q<0.05) at pH 7.0, and pH 5.7 (Fig. 2A, Supplemental Tables S2 to S4), respectively. Notably, 124 genes were induced at both pH 7.0 and pH 5.7, indicating that AC2P36 is modulating Mtb physiology independent of pH (Fig. 2B). Many of the genes upregulated at either pH are also regulated by the alternative sigma factor *sigH* in response to oxidative/thiol stress response or by hydrogen peroxide (Fig. 2ABC) (Manganelli, et al., 2002; Voskuil, et al., 2011). Indeed, *sigH* was significantly upregulated (>5-fold, q<0.05), including several genes within its regulon such as the thioredoxin reductases (*trxB, trxB2, trxC*), lipid synthesis genes (*papA4*), cysteine synthesis (*cysM*), and several predicted oxidoreductases (*Rv2454c*, *Rv3463*). Additional genes associated with sulfur-associated physiology, including sulfate transport (*cysA*, *cysW*, *cysT*) and thiosulfate sulfurtransferase (*sseC2*) are also significantly induced by AC2P36. These data are strongly indicative that AC2P36 may function by targeting thiol homeostasis. Notably, an oxidative stress transcriptional response associated with detoxification of superoxide species through superoxide dismutase (*sodA*, *sodC*) or the membrane-associated oxidoreductase (*sodA*, *sseA*, *doxX*) complex was not observed, uncoupling a general oxidative stress response from AC2P36 treatment (Nambi, et al., 2015). However, the catalase/peroxidase *katG* and ferric uptake regulator *furA* are upregulated >4-fold at both pH 7.0 and 5.7, and numerous H_2_O_2_ regulated genes are induced by AC2P36, suggesting that reactive oxygen species may be generated and require detoxification. Taken together, these data suggest that AC2P36 impacts the Mtb transcriptome at both neutral and acidic pH. However, the enhanced sensitivity of Mtb to AC2P36 treatment at acidic pH is likely due to a pH-dependent physiology, possibly via the previously described pH-dependent regulation of Mtb thiol homeostasis (Baker, et al., 2014; Buchmeier, et al., 2006; Mehta, et al., 2016).

**Figure 2.**
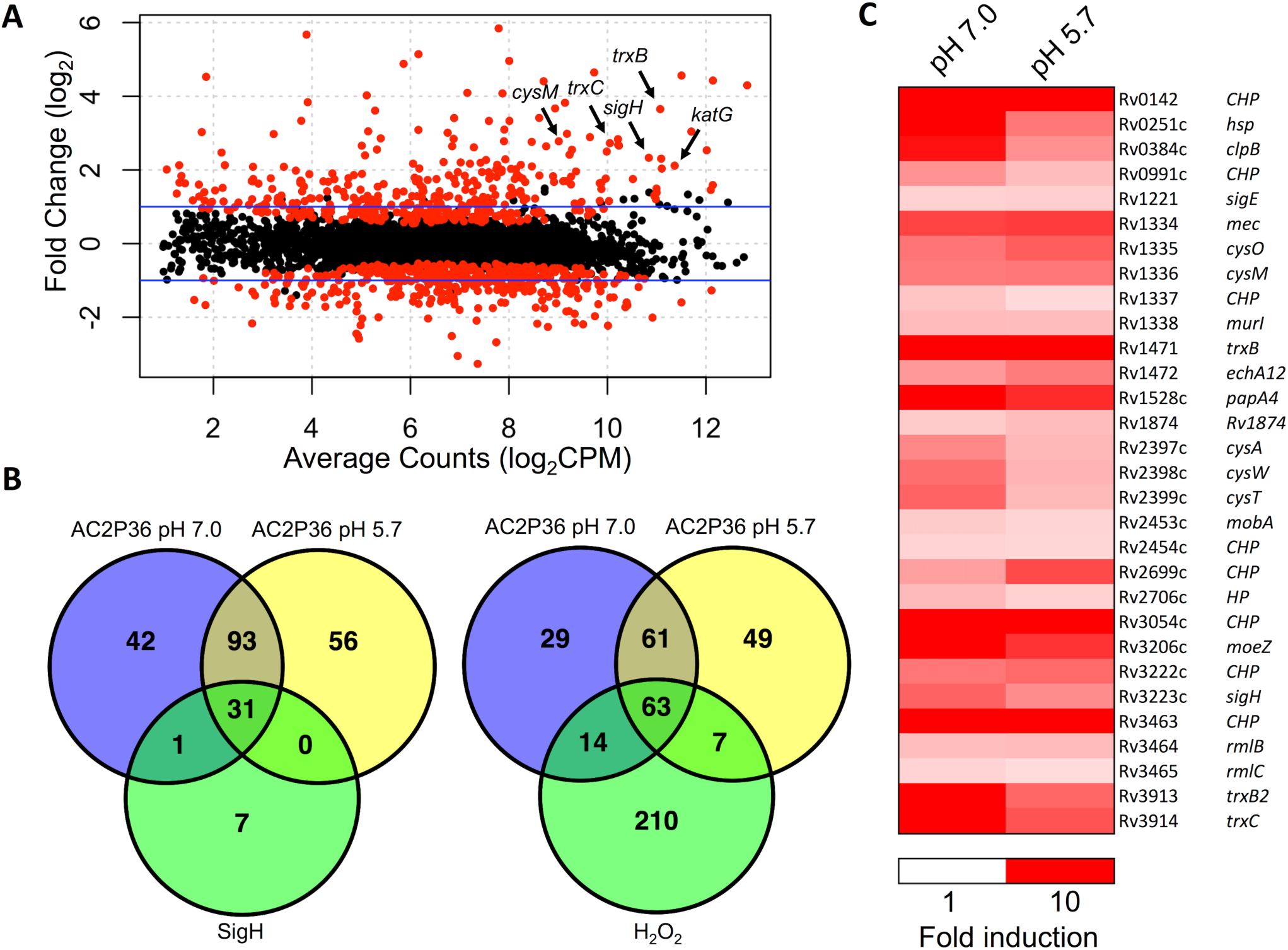
AC2P36 stimulates genes associated with the *sigH* regulon and oxidative stress. A) RNA-seq magnitude-amplitude plot of Mtb treated with 10 μM AC2P36 for 4 hours (pH 5.7). Highlighted genes are involved in the alternative sigma factor *sigH* regulon and *katG* is an Mtb catalase/peroxidase. Black dots are not statistically significant and red dots are statistically significant (q<0.05). B) Venn diagram of genes upregulated (>2-fold, q<0.05) in AC2P36 treated Mtb at pH 7.0 and 5.7 overlapping with the *sigH* regulon or H_2_O_2_ regulated genes (Manganelli, et al., 2002; Voskuil, et al., 2011). Note the significant overlap between both neutral and acidic pH in AC2P36 treated cultures. C) Heatmap showing sigH regulon genes that are induced (>2 fold, q<0.05) by AC2P36. Data are the average of two biological replicates. Comparisons are only with genes annotated in the H37Rv genome.

The transcriptional profiling data suggest that AC2P36 promotes a thiol-associated stress response, therefore, we hypothesized that AC2P36 may disrupt the bacterium’s redox poise and sensitize Mtb to oxidizing agents and antimicrobials such as diamide, clofazimine, and isoniazid (INH). To determine whether AC2P36 causes enhanced sensitivity to thiol oxidation by diamide, Mtb was inoculated into rich medium buffered at pH 5.7 in the presence or absence of 10 μM AC2P36 in combination with either 1 mM diamide or an equivalent volume of DMSO. After 3 days of treatment, cultures were plated onto 7H10 agar plates and bacterial viability was assessed through enumerating colony-forming units (CFU). Treatment with AC2P36 or diamide alone resulted in an approximately 2-fold reduction or 1.8-fold enhancement in bacterial growth relative to the initial inoculum, respectively (Fig. 3A). However, the combination of diamide and AC2P36 resulted in greater than 8-fold enhanced killing relative to the initial inoculum, demonstrating potentiating interactions of the compounds (Fig. 3A). This finding supports that AC2P36 may sensitize Mtb to thiol oxidation. Clofazimine, a component of the multidrug treatment regimen for *Mycobacterium leprae* infection, is a redox cycling compound that can directly compete with menaquinone and spontaneously produce reactive oxygen species (Lechartier and Cole, 2015). If AC2P36 functions to deplete thiol pools and diminish resistance to oxidative stress, then AC2P36 is predicted to sensitize Mtb to clofazimine. To test this hypothesis, 10 μM AC2P36 was incubated in combination with 100 μM clofazimine for 3 days at pH 5.7. Clofazimine treatment alone did not impact Mtb growth, however, clofazimine in combination with AC2P36 resulted in approximately a 22-fold reduction in bacterial growth (Fig. 3A). These data suggest that treatment with AC2P36 disrupts resistance mechanisms to oxidizing agents. Furthermore, given the induction of *katG* by AC2P36, we hypothesized that AC2P36 may sensitize Mtb to INH (a pro-drug that is activated by KatG). To test this hypothesis, Mtb was treated for 3 days in rich pH 5.7 buffered medium in the presence or absence of 10 μM AC2P36 in combination with 0.3 μM rifampin, 10 μM INH, or an equivalent volume of DMSO. AC2P36 treatment and INH alone resulted in comparable levels of killing, with approximately 2-fold reduction in bacterial growth relative to the initial inoculum (Fig. 3A). However, in combination, AC2P36 and INH synergize to reduce bacterial survival by ~100-fold (Fig. 3A). AC2P36 treatment had no significant synergy with another anti-tuberculosis drug, rifampin, at the concentration tested (Fig. 3A). Together, these data demonstrate that AC2P36 potentiates the bacteriocidal activity of oxidizing agents and specific antibiotics.

**Figure 3.**
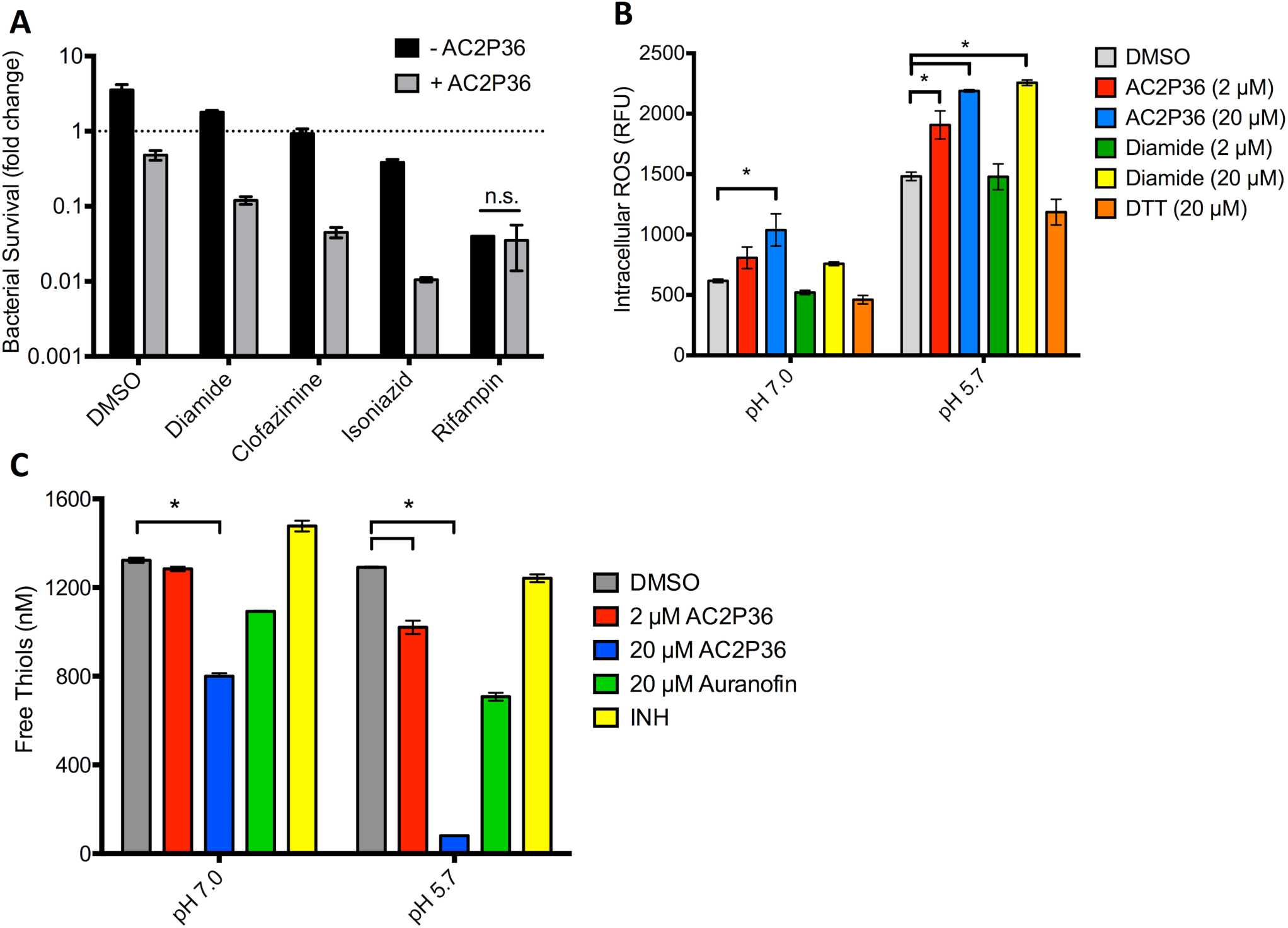
AC2P36 sensitizes Mtb to treatment by oxidizing agents, promotes the accumulation of intracellular ROS and depletes free thiols. A) Treatment of Mtb with 10 μM AC2P36 in combination with the thiol oxidizing agent diamide reduces growth ~8-fold relative to the initial inoculum. Further, the redox cycling agent clofazimine also potentiates AC2P36 treatment leading to a ~24-fold reduction in Mtb growth. Combination treatment of AC2P36 and INH results in potentiated bacterial killing (~160-fold reduction in bacterial growth). No effects were observed in combination with the antibiotic rifampin. Data are representative of two biological replicates and all differences are statistically significant (t-tests, multiple comparisons corrected p<0.05) unless indicated otherwise. n.s. - not statistically significant. B) Treatment of Mtb with AC2P36 leads to a dose-dependent and pH-dependent accumulation of reactive oxygen species (ROS). ROS was measured using the fluorescent dye CellROX Green (5 μM) and normalized by optical density (OD_595_). * represents statistical significance (p<0.05, 2-way ANOVA). C) Mtb treated with indicated concentrations of AC2P36 results in depleted thiol pools relative to the DMSO control. Auranofin is a known inhibitor that decreases the concentrations of free thiols while isoniazid (INH) is an anti-tuberculosis antibiotic that does not directly deplete free thiols. Statistical significance was calculated based on a two-tailed t-test (*, p<0.05). For each panel, the error bars represent the standard deviation of the mean.

### AC2P36 causes accumulation of intracellular reactive oxygen species

The transcriptional profiling data and potentiation of oxidation agents suggests that AC2P36 may promote the accumulation of intracellular reactive oxygen species (ROS). Recent studies have shown that increases in endogenous ROS can have significant and detrimental results on Mtb growth and viability *in vitro* and during infection (Vilcheze, et al., 2013) (Tyagi, et al., 2015). To measure intracellular ROS formation in Mtb, the fluorescent dye CellROX green was employed (Saini, et al., 2016). Mtb was grown in rich medium, pelleted, and re-suspended in either pH 7.0 or 5.7 buffered 7H9 medium. Cultures were incubated in the presence of 2 or 20 μM AC2P36 or an equivalent volume of DMSO. At both acidic and neutral pH, AC2P36 caused a significant induction of ROS (Fig. 3B). AC2P36 in combination with acidic pH doubles the accumulation of intracellular ROS as compared to AC2P36 treated cells at neutral pH. Notably, acidic pH alone causes an accumulation of ROS, suggesting that acidic pH stress causes enhanced ROS formation or inhibits oxidative stress resistance mechanisms. To test the hypothesis that acidic pH reduces the ability of Mtb to counteract oxidative stress, we examined intracellular ROS accumulation in diamide treated cells at acidic pH. Diamide at a concentration of 20 μM causes a significant, pH-dependent increase in ROS at pH 5.7. These data are consistent with Mtb having diminished resistance to oxidative stress at acidic pH.

### AC2P36 depletes intracellular thiol pools

We hypothesized that AC2P36 may deplete thiol pools in an acidic pH-dependent manner, given that AC2P36 promotes thiol stress and sensitivity to oxidizing agents, and that acidic pH has been shown to promote reductive stress and enhanced accumulation of the small molecular weight thiol buffer mycothiol (Buchmeier, et al., 2006). To directly measure intracellular free thiol pools, Mtb was inoculated into rich medium buffered at pH 7.0 and pH 5.7 in the presence of either 2 μM AC2P36, 20 μM AC2P36, 20 μM auranofin, INH, or an equivalent volume of DMSO. Following 24 hours incubation, cultures were analyzed for total intracellular thiol pools. Previous work has shown that auranofin depletes free thiols in Mtb and *S. aureus;* therefore it was used as a positive control (Harbut, et al., 2015). INH was used as a negative control as it is not anticipated to directly deplete free thiols. Indeed, auranofin depleted free thiols relative to the DMSO treated control and INH had comparable thiol concentrations as the untreated cultures (Fig. 3C). AC2P36 significantly depleted thiol pools at both pH 7.0 and 5.7. At pH 5.7, 20 μM AC2P36 depleted free thiols ~16 fold to ~80 nM as compared to ~1.3 μM in the DMSO treated samples (Fig. 3C), whereas at pH 7.0 free thiols were only reduced by one third. Overall, these data show that AC2P36, i) promotes a reduced cytoplasm, ii) increases intracellular ROS, and iii) significantly depletes free thiol pools, with enhanced depletion at acidic pH. These data demonstrate that Mtb is more sensitive to thiol targeting agents at acidic pH and that this pH-dependent sensitivity enhances vulnerability to oxidizing agents and antibiotics. The findings are consistent with prior reports that Mtb or *M. smegmatis* mutants deficient in mycothiol are highly sensitive to redox-cycling agents, H_2_O_2_ or ascorbic acid (vitamin C) (Buchmeier, et al., 2006; Rawat, et al., 2002; Vilcheze, et al., 2013). Interestingly, mycothiol mutants have been shown to have enhanced resistance to INH (Vilcheze, et al., 2011; Xu, et al., 2011) (Rawat, et al., 2002), in contrast to the INH potentiation caused by AC2P36, indicating that the AC2P36 mechanism of action is not exclusively interfering with the mycothiol redox cycling pathway.

### AC2P36 modulates *S. aureus* physiology with mechanisms similar to Mtb

Given that AC2P36 can also inhibit *S. aureus* growth, we examined if AC2P36 similarly increased intracellular ROS and depleted free thiols *S. aureus.* To examine intracellular ROS, *S. aureus* was inoculated into rich medium buffered to either pH 7.0 or 5.7 and exposed to a 4 point (2-fold) dilution series of AC2P36, DTT, and menadione ranging from 10 - 80 μM or an equivalent volume of DMSO. Following 6 hours incubation, cultures were assessed for accumulated ROS. AC2P36 caused enhanced ROS accumulation in a dose-dependent manner at pH 7.0 and 5.7 similar to the positive menadione control (Fig. 4A). Notably, ROS accumulated at higher levels at neutral pH as compared to acidic pH, indicating differences exist in pH-dependent redox homeostasis between Mtb and *S. aureus.* AC2P36 also significantly reduced free thiols in *S. aureus,* with both 20 μM AC2P36 or auranofin causing a ~30% reduction in free thiols (Fig. 4B). The enhanced accumulation of ROS may be directly driving growth inhibition. To test this hypothesis, AC2P36 sensitivity was tested with the superoxide dismutase deficient *S. aureus* (*ΔsodAΔsodM*) mutant, which has reduced resistance to oxidative stress (Kehl-Fie, et al., 2011). As compared to the WT, *S. aureus* (*ΔsodAΔsodM*) exhibited a ~2-fold enhanced sensitivity to AC2P36 at pH 7.0 and 5.7 (Fig. 4C). These data suggest that AC2P36 treatment results in accumulated intracellular ROS in *S. aureus* and that growth inhibition is driven, in part, by an ROS mediated mechanism.

**Figure 4.**
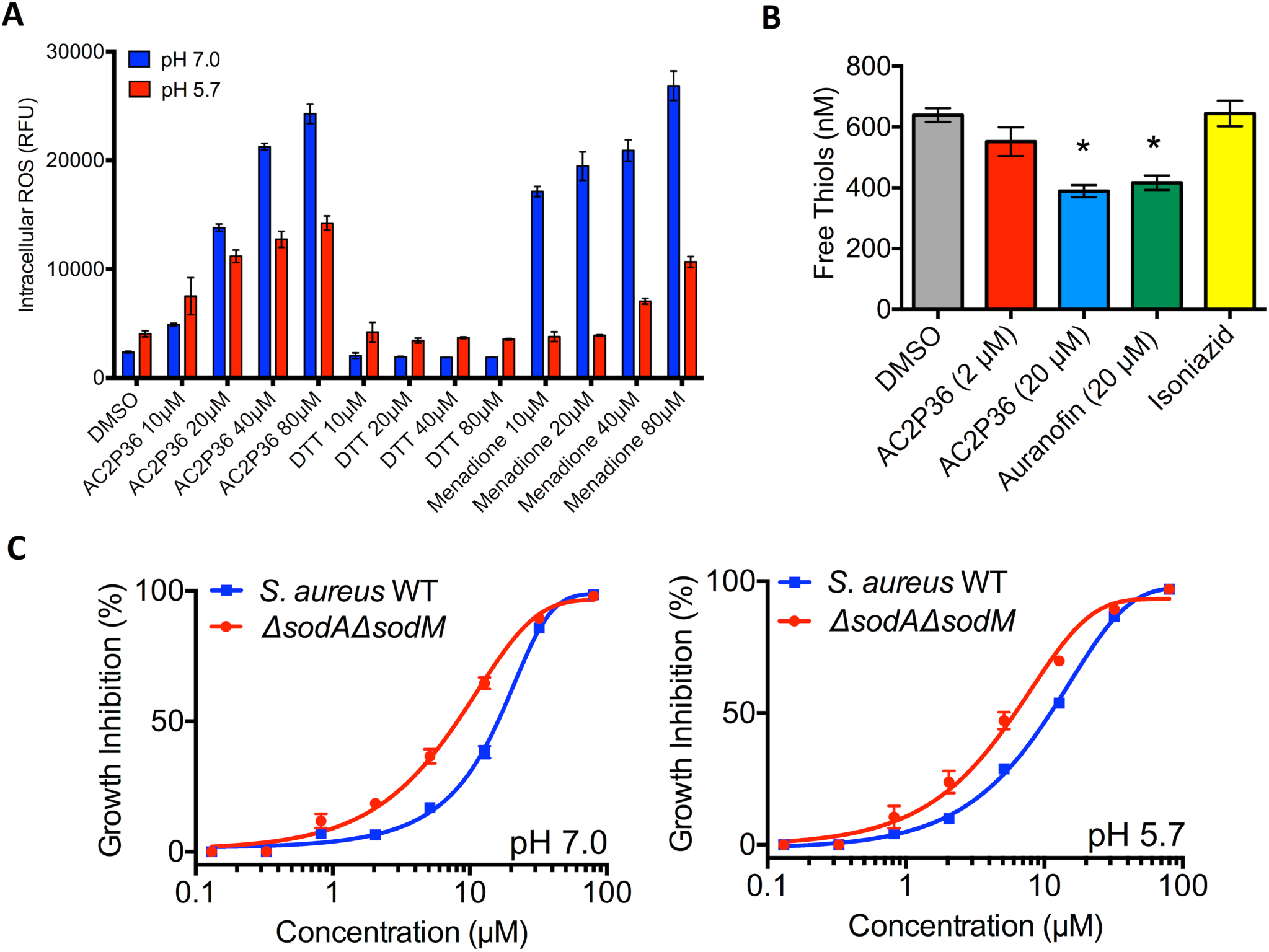
AC2P36 stimulates ROS production and depletes free thiols in *S. aureus*. A)Treatment of S. *aureus* with indicated concentrations of AC2P36 leads to a dose-dependent accumulation of reactive oxygen species (ROS). In contrast to Mtb, *S. aureus* exhibits enhanced ROS accumulation at neutral pH. *S. aureus* treated with AC2P36 exhibits a similar dosedependent response to the redox cycling agent menadione. DTT had no effect on ROS accumulation. ROS was measured using the fluorescent dye CellROX Green (5 μM) and normalized by optical density (OD_595_). All differences in *S. aureus* are statistically significant between the DMSO treated control and AC2P36 treated cells (p<0.05, 2-way ANOVA). B) *S.aureus* treated with indicated concentrations of AC2P36 at pH 5.7 results in depleted thiol pools relative to the DMSO control. Statistical significance was calculated based on a two-tailed t-test (*, p<0.05). C) *S. aureus* sensitivity to AC2P36 is enhanced ~ 2 fold in the *S. aureus (AsodAAsodM)* mutant at both pH 7.0 and pH 5.7, supporting that growth inhibition is partially driven by oxidative stress.

### The AC2P36 sulfone group is required for activity

Eukaryotic cytotoxicity of AC2P36 was tested in J774 macrophages and it was observed that the compound had a half-maximal cytotoxicity concentration (CC_50_) of 17 μM. Therefore, we sought to define the chemical groups required for activity against Mtb while minimizing cytotoxic effects against mammalian cells. To define the reactive moieties of AC2P36, the core chemical structure was determined and several analogs of the parent molecule (AC2P36-A, Fig. 5) were assessed for bactericidal activity or eukaryotic cytotoxicity (Fig. 5). Changes in the chemical groups at the R1 and R2 regions of the core structure revealed that the sulfone moiety at the R1 position is required for activity against Mtb at pH 5.7. In the AC2P36-H and AC2P36-I analogs, the methylsulfone group has been replaced with alternative groups, rendering both molecules inactive against Mtb. Notably, the AC2P36-J analog reveals that the R2 moiety is not required for killing of Mtb. Further, this AC2P36-J analog does not exhibit cytotoxicity against macrophages up to the highest concentration tested (80 μM), suggesting that the 5-chloro-N-(3-chloro-4-methoxyphenyl) R2 group may drive the cytotoxicity of the parent compound; although, it cannot be ruled out that loss of the R2 moiety may limit entry of the AC2P36-J analog into the macrophage. These data suggest that the reactive group of AC2P36 resides in the methylsulfonylpyrimidine scaffold and that the sulfone moiety is required for activity.

**Figure 5.**
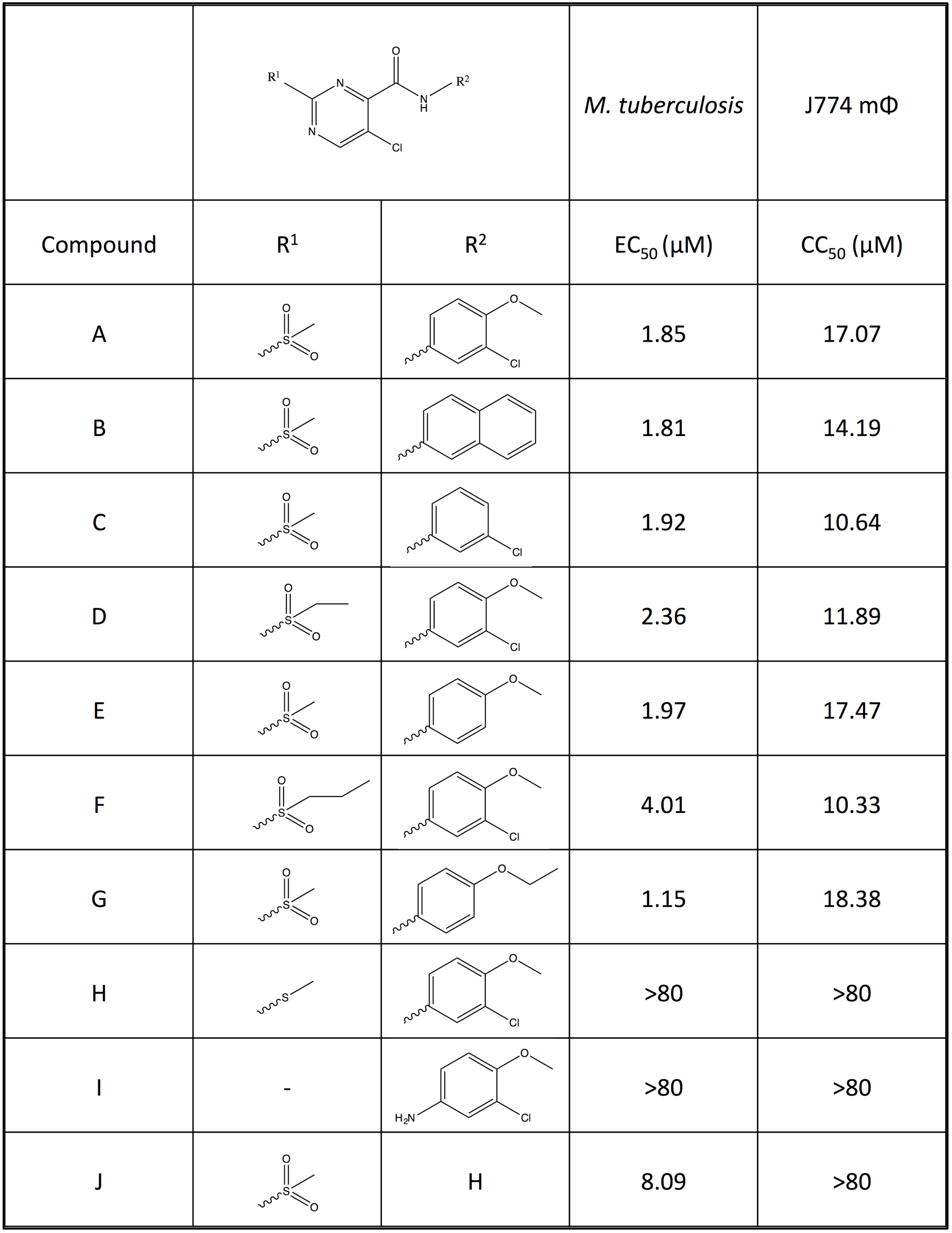
Structure activity relationship studies of AC2P36. Testing of growth inhibition and macrophage cytotoxicity (CC_50_) of the parent analog AC2P36-A and 9 additional analogs (B-J) with variations at the specified R1 and R2 groups. The R2 group is dispensable, while the R1 group is indispensible for inhibition of Mtb growth at pH 5.7. Notably, eukaryotic cytotoxicity is abolished in the Mtb active analog AC2P36-J.

### AC2P36 directly targets thiols

The core chemical scaffold of AC2P36 (5-chloro-2-methylsulfonylpyrimidine-4-carboxamide) has been researched previously for synthesizing derivative scaffolds of aminopyrimidine-containing drugs (Moon, et al., 2013). Notably, the anionic nucleophile thiolate was shown to selectively react with the C-2 carbon almost exclusively depending on the electron-withdrawing group (Moon, et al., 2013). In the case of a methylsulfone moiety, this chemical group is a strong electron-withdrawing group, promoting a partial positive charge at the C-2 carbon allowing the electron rich thiolate anion to attack and displace the methylsulfone (Bebbington, et al., 2009). Based on the known chemistry, we hypothesized that AC2P36 may directly target thiols. To test this hypothesis, we incubated 80 μM AC2P36 with 100 μM reduced glutathione (GSH) or a DMSO control at both acidic and neutral conditions for one hour. The samples were then examined by mass spectrometry. In the AC2P36 alone treated samples, AC2P36 was detected as a peak with a mass of 373.976 Da. However, in the AC2P36 and GSH treated samples at both acidic and neutral pH, a new peak was observed at 601.066 Da (Fig. 6A). This new peak is consistent with the proposed reaction shown in Figure 6B, where the covalent modification of free thiols by AC2P36 promotes the release of either methanesulfinic acid or methanesulfinate. For example, the combined masses of AC2P36 (373.976 Da) and GSH (307.32 Da) is 681.246 Da and the loss of methanesulfinic acid (79.065 Da) and a proton, results in a molecule with a mass of 601 Da. These data support that AC2P36 is capable of forming stable adducts with free thiols and support the proposed reaction mechanism.

**Figure 6.**
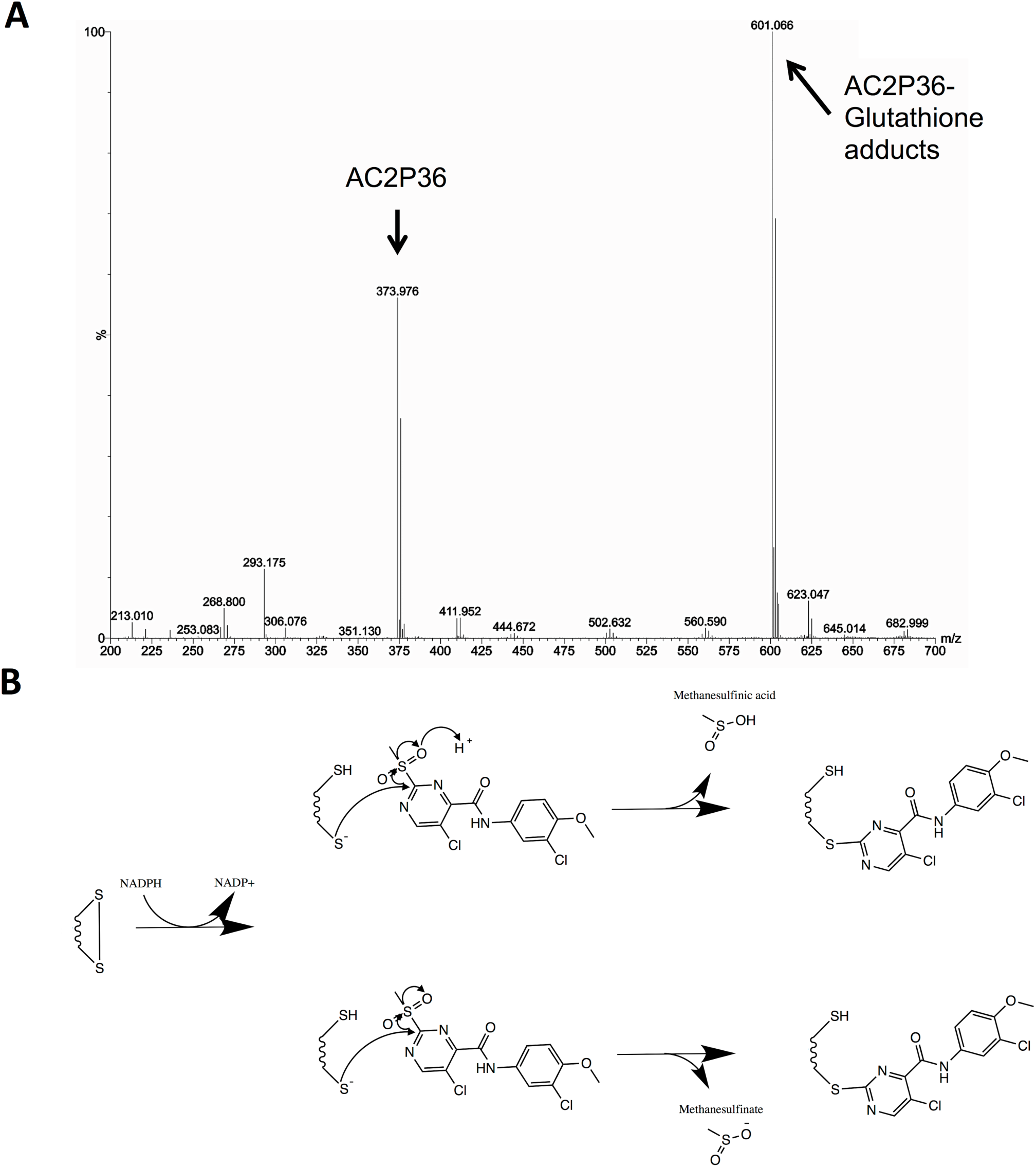
AC2P36 forms covalent adducts with free thiols. A) Incubation of AC2P36 with reduced glutathione (GSH) results in the formation of AC2P36-GSH adducts as evidenced by formation of a new molecule with a molecular weight of ~601 Da. B) A proposed reaction mechanism that could generate the observed AC2P36-GSH adduct. NADPH-dependent activation of the general disulfide to a thiol and thiolate anion allows for the thiolate to undergo anionic nucleophilic attack of the C-2 carbon on the pyrimidine ring of AC2P36, leading to a covalent modification and releasing either methanesulfinic acid or methansulfinite.

## Discussion

AC2P36 was discovered as a pH-dependent inhibitor of Mtb growth by conducting two whole-cell HTS of an identical compound library at neutral and acidic pH. Mechanism of action studies support that that AC2P36 depletes intracellular thiol pools and promotes intracellular accumulation of ROS and these activities may lead to the observed bacteriocidal activity against Mtb. Notably, Mtb exhibits reduced thiol pools, ROS accumulation and associated changes in gene expression at both acidic and neutral pH, supporting that AC2P36 is modulating Mtb physiology independently of pH. However, the magnitude of the perturbations is enhanced at acidic pH, and Mtb is selectively killed by AC2P36 at acidic pH. Therefore, we propose that Mtb has enhanced sensitivity to AC2P36-driven thiol stress at acidic pH. These observations are supported by recent studies showing that acidic pH promotes metabolic stress associated with a reduced cytoplasm (Baker, et al., 2014; Mehta, et al., 2016). A previous study showed that mycothiol pools increase approximately two-fold at acidic pH (pH 5.5), suggesting Mtb is adapting to thiol stress at acidic pH (Buchmeier, et al., 2006). Notably, the thiol pool assays performed in this study (Fig. 3C) did not detect changes in thiol pools at acidic versus neutral pH in the DMSO treated controls. However, several differences exist between the studies, including the duration of acid stress and sensitivity of the thiol detection assays (Buchmeier, et al., 2006).

Given that Mtb experiences a reduced cytoplasm at acidic pH, we propose a model where Mtb enhanced sensitivity to AC2P36 at pH 5.7 is the result of a greater proportion of thiol pools being in a reduced state. AC2P36 could then covalently modify the reduced thiols, thus inhibiting their ability to be re-oxidized (Fig. 7). This model mechanism is supported by the redox-sensitive GFP data indicating a reduced cytoplasm following AC2P36 treatment (Fig. 1E), the transcriptional profiling results indicating thiol and pH-dependent AC2P36-mediated enhanced reductive stress, and greater depletion of thiol pools at acidic pH relative to neutral pH (Fig. 3C). Given that we did not observe enhanced levels of free thiols at acidic pH, it is possible that the reduced intracellular environment is sufficient to maintain thiol pools in a reduced and thus, an AC2P36 susceptible state. Thiol recycling pathways play an important role in counteracting oxidative stress, therefore, it follows that AC2P36 may inhibit the ability of Mtb to dissipate oxidative stress and thus lead to the observed enhanced accumulation of ROS and impaired resistance to oxidizing agents. We hypothesize that insensitivity of Mtb to AC2P36 at neutral pH is driven by the more oxidized cytoplasm, resulting in AC2P36-resistant oxidized thiols, or alternatively, auxiliary ROS resistance or thiol recycling pathways may be available to the bacterium at neutral pH. Mtb is predicted to occupy acidic niches during infection, therefore, our findings provide a proof of concept study showing that targeting free thiol pools is a potential therapeutic strategy for Mtb drug development.

**Figure 7.**
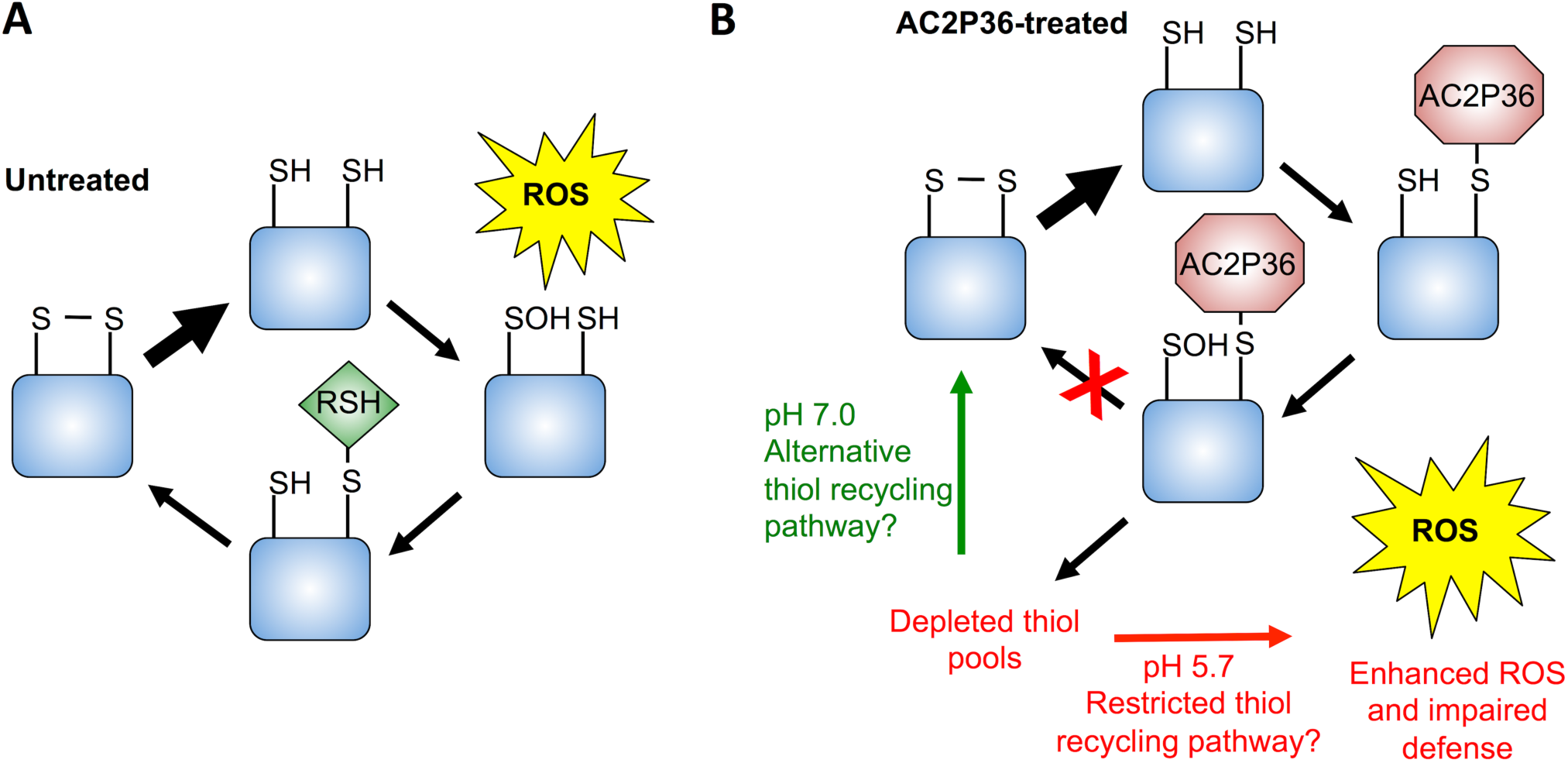
Model mechanism for the AC2P36-dependent inhibition of growth and potentiation of oxidizing agents. A) Normal responses of Mtb to reactive oxygen species (ROS) stress at neutral and acidic pH can proceed through recycling of oxidized thiol pools via a low molecular weight thiol (RSH) and mycoredoxin/thioredoxin (not shown) to regenerate a disulfide bond. B) At neutral pH, AC2P36 depletes thiol pools but Mtb possesses the ability to maintain or regenerate thiol pools to defend against ROS. However, at acidic pH, Mtb has a reduced cytoplasm resulting in an enhanced ratio of reduced thiols to oxidized disulfides. Treatment with AC2P36 covalently modifies free thiols preventing Mtb from maintaining or regenerating intracellular thiol pools and redox buffering. As a result, Mtb is unable to defend against oxidative damage, leading to cell death and potentiation of oxidizing agents.

AC2P36 strongly induces many genes within the alternative sigma factor SigH regulon, suggesting that Mtb experiences thiol stress when treated with AC2P36 (Manganelli, et al., 2002). The SigH regulon was identified based on comparing the thiol oxidizing agent diamide compared to a diamide treated *sigH* mutant. Induced genes in the SigH regulon include thioredoxins, thioredoxin reductase, as well as genes associated with cysteine biosynthesis. Thioredoxins and the thioredoxin reductase are key enzymes in the process of maintaining redox homeostasis in Mtb. If the ability to recycle thiols through the thioredoxin reductase is inhibited, Mtb is attenuated for growth (Harbut, et al., 2015). However, AC2P36 likely does not directly inhibit thioredoxin reductase given the enzyme is essential for growth independent of pH (Harbut, et al., 2015). The observed upregulation of the thioredoxins and thioredoxin reductase at both pH 7.0 and 5.7 suggests that AC2P36 is potentially blocking the ability of Mtb to recycle its thiol pool to maintain redox balance. Additionally, the upregulation of cysteine biosynthesis has been implicated in maintaining redox balance in Mtb strains deficient in the low molecular weight thiol ergothioneine and the redox sensor WhiB3 (Saini, et al., 2016). However, we did not observe upregulation of either mycothiol or ergothioneine biosynthetic operons by AC2P36 at either neutral or acidic pH, suggesting that AC2P36 activity does not directly promote increased production of low molecular weight thiols as a countermeasure. The upregulation of the multifunctional catalase/peroxidase KatG, may be explained in part by oxidation of NADH by KatG to maintain redox balance and counteract AC2P36 activity (Singh, et al., 2004). However, this function of KatG is also potentially detrimental to the cell as hydrogen peroxide is produced in the process of consuming reducing equivalents (Farhana, et al., 2010; Singh, et al., 2004). We observe both enhanced accumulation of ROS and a ~100-fold potentiation of INH when Mtb is treated with AC2P36 and thus propose that induction of KatG may drive the enhanced susceptibility to the pro-drug INH, although additional mechanisms likely drive the synergistic interaction. Notably, it was previously shown that modulation of thiol homeostasis, through manipulation of mycothiol synthesis, causes INH resistance (Rawat, et al., 2002; Vilcheze, et al., 2011; Xu, et al., 2011), therefore, targeting of thiol homeostasis may enhance or reduce INH potency depending on the mechanism of the inhibition.

The proposed mechanism for AC2P36 suggests that the compound is generally thiol-reactive and would also target eukaryotic thiol pools. This possibility is highlighted by the cytotoxic effects of AC2P36 against eukaryotic macrophages (Fig. 5). However, the AC2P36-J analog remains active against Mtb while showing no macrophage cytotoxicity, suggesting that Mtb specific functions may exist for this compound. Further, in light of the known chemistry surrounding this scaffold, it may be possible to develop inhibitors of Mtb thiol pools that act as a pro-drug, resulting in activation of the thiol targeting chemistry only inside the bacterium. Indeed, the nitric oxide producing drug PA-824 is a pro-drug that specifically promotes nitrosative stress inside the bacterium (Singh, et al., 2008).

## Significance

*Mycobacterium tuberculosis* (Mtb) must adapt to acidic environments to colonize and survive in the host. Therefore, discovery of pathways that are required for survival at acidic pH and vulnerable to chemical inhibition may identify novel antivirulence approaches to control Mtb infection. Here we describe a small molecule, AC2P36, which selectively kills Mtb at acidic pH. AC2P36 is shown to potently deplete free thiol pools and promote the accumulation of reactive oxygen species at acidic pH. AC2P36 strongly synergizes with the first line antibiotic isoniazid, supporting that modulating Mtb free thiol pools may enhance the efficacy of Mtb treatment. These data support a model where chemically depleting Mtb thiol pools at acidic pH leads to enhanced sensitivity to oxidative damage, resulting in bacterial killing and potentiation of antibiotics.

## Experimental Procedures

### Bacterial strains and growth conditions

*Mycobacterium tuberculosis* (Mtb) experiments, unless otherwise stated, were performed with Mtb strain CDC1551. Cultures were maintained in standing tissue culture flasks in 7H9 Middlebrook medium supplemented with 10% oleic acid, albumin, dextrose, catalase (OADC) and 0.05% Tween-80 and incubated at 37^o^C with 5% CO_2_ unless noted otherwise. *Staphylococcus aureus* experiments were performed with *S. aureus* strain Newman and maintained in shaking 15 mL Falcon tubes or Erlenmeyer flasks in Luria-Bertani (LB) broth. Where necessary, 7H9 or LB media were buffered to pH 7.0 with 100 mM MOPS or pH 5.7 with 100 mM MES (Piddington, et al., 2000).

### EC_50_ determination and spectrum of activity in non-mycobacteria

For EC_50_ determinations, cultures were incubated in 96-well microtiter plates in the presence of an 8-point (2.5-fold) dilution series of AC2P36 ranging from 80 μM – 0.13 μM. Mtb cultures were grown to mid- to late-log phase and then pelleted, re-suspended in 7H9 buffered at pH 7.0 or 5.7 and dispensed into the 96-well assay plates at an optical density (OD) of 0.1 and incubated for 6 days at 37°C with 5% CO_2_. The activity of AC2P36 against non-mycobacteria was performed in buffered pH 7.0 or pH 5.7 LB broth, shaking at 200 RPM at 37°C, except for *Enterococcus faecalis,* which was grown in Brain Heart Infusion medium in standing cultures at flasks at 37°C. Cultures were grown overnight and then back diluted to a starting optical density (OD) of 0.05 in 96-well microtiter plates. Several Gram-positive and Gram-negative bacteria were tested including: *Staphylococcus aureus* strains Wichita (29213) or Seattle (25923), *Escherichia coli* (Migula) Castellani and Chalmers, *Pseudomonas aeruginosa* (Schroeter)Migula and, *Proteus vulgaris* Hauser emend. Judicial Commission, *Enterococcus faecalis* (Andrewes and Horder) Schleifer and Kilpper-Balz. Bacteria were incubated in the presence of an 8-point (2.5-fold) dilution series of AC2P36 ranging from 80 μM – 0.13 μM for a period of 6-8 hours. Growth (OD) was monitored using a Perkin Elmer EnVision plate reader. Non-mycobacteria cultures were normalized for growth based on the kanamycin (100%) and DMSO (0%) controls, with the exception of *P. aeruginosa,* where tobramycin was used as the 100% positive control. EC_50_ values were calculated based on a variable slope, four-parameter nonlinear least squares regression model in the GraphPad Prism software package (ver. 6).

### Bactericidal Activity of AC2P36

Mtb cultures were prepared at an OD_600_ of 0.1 in buffered 7H9 media (pH 5.7) in 96-well plates as described above. Mtb cultures were treated with AC2P36 (at indicated concentrations) for a period of 3 days. At the end of the incubation, 10-fold serial dilutions were performed for each condition and viable bacterial CFUs were enumerated by plating on 7H10 agar. Bactericidal activity was determined by comparing bacterial CFUs after treatment to the initial inoculum. Cultures treated with DMSO and rifampin were included as negative and positive controls, respectively.

### Measuring intracellular redox state

The intracellular redox state of Mtb cultures treated with the compounds of interest was determined using the redox-sensitive Mtb reporter strain CDC1551⸬roGFP-R12 (Baker, et al., 2014; Cannon and Remington, 2006; Hanson, et al., 2004). Briefly, Mtb was cultured in buffered pH 7.0 or 5.7 7H9 media lacking catalase (e.g. OAD) in 96-well plates containing 0.1 – 20 μM AC2P36 at a starting OD_600_ of 0.2. For each experimental condition, non-fluorescent wild-type CDC1551 transformed with empty vector was used as a control for background signal subtraction. At 6 h post-treatment, fluorescence emission was read at 510 nm after excitation at 400 nm and 480 nm, and the relative abundance of the oxidized and reduced roGFP species were measured, respectively. The 400/480 nm ratio of drug-treated cultures were normalized to the DMSO control which represented the baseline redox state of the cell. Dithiothreitol (DTT) was used as a control for a reductive environment and diamide was used as a control for oxidizing environments.

### Transcriptional profiling and data analysis

For RNA-seq experiments, Mtb cultures were grown at 37°C in T-25 vented, standing tissue culture flasks in 8 mL of buffered 7H9 medium (pH 7.0 or 5.7) to an OD_600_ of 0.5. Cultures were subsequently treated with 10 μM AC2P36. As a baseline control, cultures treated with an equal amount of DMSO were used. Each treatment was repeated with two independent (biological) replicates. Following a 4 h incubation period in the presence of AC2P36, total bacterial RNA was extracted, quality controlled and sequenced as previously described (Baker, et al., 2014; Rohde, et al., 2007). RNA-seq data was analyzed using the SPARTA software package (Johnson, et al., 2016). Genes with an average log_2_CPM <5 were filtered from the final differential gene expression lists (Supplemental Tables S2AB, S3AB, S4AB). Genes with >2 fold induction or repression and a false discovery rate corrected p-value (denoted q) <0.05, were included in the lists of significantly differentially regulated genes. The transcriptional profiling data is available at the NCBI GEO database (GSE89106).

### Structure activity relationship (SAR) and eukaryotic cytotoxicity of AC2P36

Several analogs of AC2P36 were purchased to determine the structure activity relationships. Compounds B-J (Table 5.2.) were assessed for growth inhibitory activity in Mtb using an 8-point (2.5-fold) dilution series ranging from 80 – 0.13 μM at pH 5.7 in a similar manner as EC_50_ determinations described above. Following 6 days incubation, cultures were analyzed for growth using OD relative to the rifampin (100%) and DMSO (0%), positive and negative controls, respectively.

Eukaryotic cytotoxicity of AC2P36 and its analogs were assessed as previously described (Johnson and Abramovitch, 2015). Briefly, J774 macrophages were grown at 37°C (5% CO_2_) in DMEM (Corning CellGro) containing 10% fetal bovine serum (Thermo Scientific), 1 mM pyruvate, 2 mM L-glutamine, and 1 mM penicillin/streptomycin (Corning CellGro). Cells were grown until confluent, scraped, and plated in medium lacking antibiotics at either 1 x 10^5^ cells/mL. Cells were allowed to adhere overnight before addition of experimental treatments. Cells were incubated for 3 days at 37°C with 5% CO_2_ and assessed for viability using the CellTiter Glo (Promega) luciferase kit. Cytotoxicity was normalized based on the 1% Triton X-100 (100%) and DMSO (0%) controls. EC_50_ and half-maximal cell cytotoxicity concentration (CC_50_) values were calculated based on a variable slope, four-parameter non-linear least squares regression model in the GraphPad Prism software package (ver. 6).

### Fractional killing and potentiation of antimicrobials

Mtb cultures were grown to mid- to late-log phase, pelleted, and re-suspended in buffered pH 7.0 or 5.7 7H9 lacking catalase at an OD of 0.1. Cells were assessed for viability in the presence or absence of 10 μM AC2P36 in combination with: DMSO, 1 mM diamide, 100 μM clofazimine, 0.3 μM rifampin, or 10 μM isoniazid. Following 3 days incubation in 96-well microtiter plates, cells were diluted in a 10-fold serial dilution series for each condition and viable bacterial CFUs were enumerated by plating on 7H10 agar.

### Measurement of intracellular reactive oxygen species and pH

Intracellular reactive oxygen species were measured using the CellROX Green reagent (Invitrogen) as previously described (Saini, et al., 2016). Briefly, Mtb was grown to mid- to late-log in Middlebrook 7H9 and inoculated into 5 mL of buffered 7H9 pH 7.0 or 5.7 lacking catalase at a starting OD of 0.5 and incubated in the presence of indicated concentrations of compounds. Following 4 or 24 hours incubation, a final concentration of 5 μM CellROX green (Thermo Fisher) was added to cultures for 1 hour at 37 °C. Cells were washed twice in PBS supplemented with 0.05% Tween80 and re-suspended in the wash buffer and analyzed according to the manufacturers instructions. DTT (20 μM), and diamide (2 and 20 μM) served as controls. *S. aureus* cultures were grown overnight in LB at 37°C with shaking at 200 RPM. Cultures were pelleted and re-suspended in buffered pH 7.0 or pH 5.7 LB medium at an OD of 0.1. Cells were treated with a 4-point (2-fold) dilution series of AC2P36, DTT, or menadione ranging from 80 – 10 μM. An equivalent volume of DMSO was added as a negative control. Following 6 hours of treatment, cultures were incubated with 5 μM CellROX Green for 30 minutes at 37°C with shaking at 200 RPM. Cells were washed twice with PBS and resuspended in 0.6 mL of PBS before being arrayed in a 96-well microtiter plate. Fluorescence and growth were analyzed using a Perkin Elmer plate reader. Cytoplasmic pH was measured over the course of 24 hours following treatment with AC2P36 at pH 5.7 as compared to DMSO or Nigericin negative and positive controls, respectively, using the method described by Purdy, Niederweis, and Russell (Purdy, et al., 2009).

### Measurement of free thiol pools

Mtb cultures were grown to mid- to late-log phase in 7H9 medium lacking catalase, pelleted, and re-suspended in 8 mL buffered pH 7.0 or 5.7 7H9 at an OD of 0.25. Five treatments were assessed: 1) DMSO, 2) 2 μM AC2P36, 3) 20 μM AC2P36, 4) 20 μM auranofin, and 5) isoniazid. Following 24 hours of treatment, bacteria were normalized by OD, pelleted, washed twice in PBS + 0.05% tyloxapol, and re-suspended in 0.75 mL thiol assay buffer (100 mM potassium phosphate pH 7.4, and 1 mM EDTA). Cells were lysed by bead beating for 2 minutes at room temperature. Supernatants were removed and assayed using the Cayman Thiol Detection Assay kit (Cayman Chemical).

### AC2P36-GSH adduct formation assay

In 100 μL Tris-HCl buffer (pH 7.0 or 5.7), 80 μM AC2P36 was added together with 100 μM reduced glutathione. The control without reduced glutathione was also performed. The reactions were incubated at RT for 1 hour. For mass spectroscopy (MS) analysis, the samples were diluted 1:100 with water, and analyzed in Waters Xevo G2-XS QTof mass spectrometer (Milford, MA, USA) with an electrospray ionization negative mode. The instrument parameters were as followed: capillary voltage, 2 kV; sampling cone, 40 V; source temperature, 100 °C; desolvation temperature, 350 °C; cone gas flow, 25 L/h; desolvation gas flow, 600 L/h. Chromatographic separation was performed in a Waters ultra-performance liquid chromatograph (ACQUITY UPLC) system. Solvents were (A) water, and (B) acetonitrile. The flow rate was 0.2 mL/min with the gradient of A/B = 50/50 for 2 min. The acquisition mass range was 50–1,500 Da. The experiment was repeated with at least two biological replicates with similar results.

## Author Contributions

GBC, BKJ, NDH and RBA conceived the experiments; GBC, BKJ, CJC, RJF, HZ and EH conducted experiments; GBC, BKJ and RBA wrote the manuscript.

## Accession Codes

The transcriptional profiling data have been submitted to the NCBI GEO database (accession no. GSE89106).

## Acknowledgements

We thank the New England Regional Center of Excellence (U54 AI057159) for providing the screening libraries and Su Chiang and Doug Flood for assistance in preparing the compound libraries for screening. The High Performance Computing Cluster and iCER at Michigan State University provided computational support. The MSU RTSF provided technical support for the RNA-seq library preparation and sequencing. We thank Eric Skaar for sharing *S. aureus* mutant strains, Javiera Ortiz for technical assistance and members of the Abramovitch lab for critical reading of the manuscript. Research reported in this study from the Abramovitch lab was supported by start-up funding from Michigan State University and AgBioResearch, and grants from the NIH-NIAID (U54AI057153 and R01AI116605) and the Jean P. Schultz Endowed Biomedical Research Fund at the MSU College of Human Medicine.

---

**Supplemental Figure 1.**
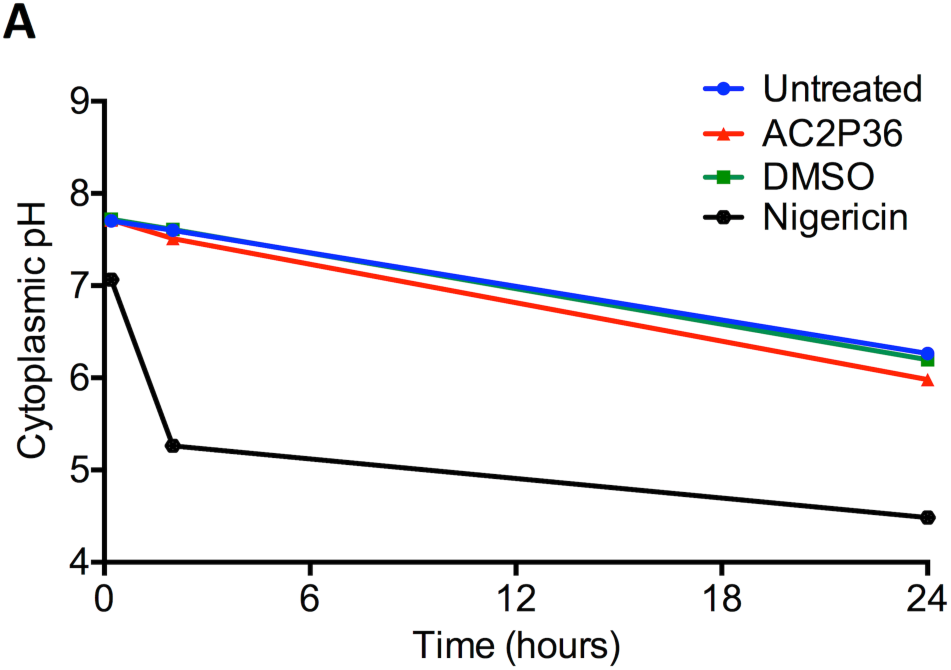
AC2P36 does not impact the cytoplasmic pH of Mtb at pH 5.7. DMSO and nigericin are added as negative and positive controls. Data are representative of two independent experiments.

**Supplemental Table 1.**
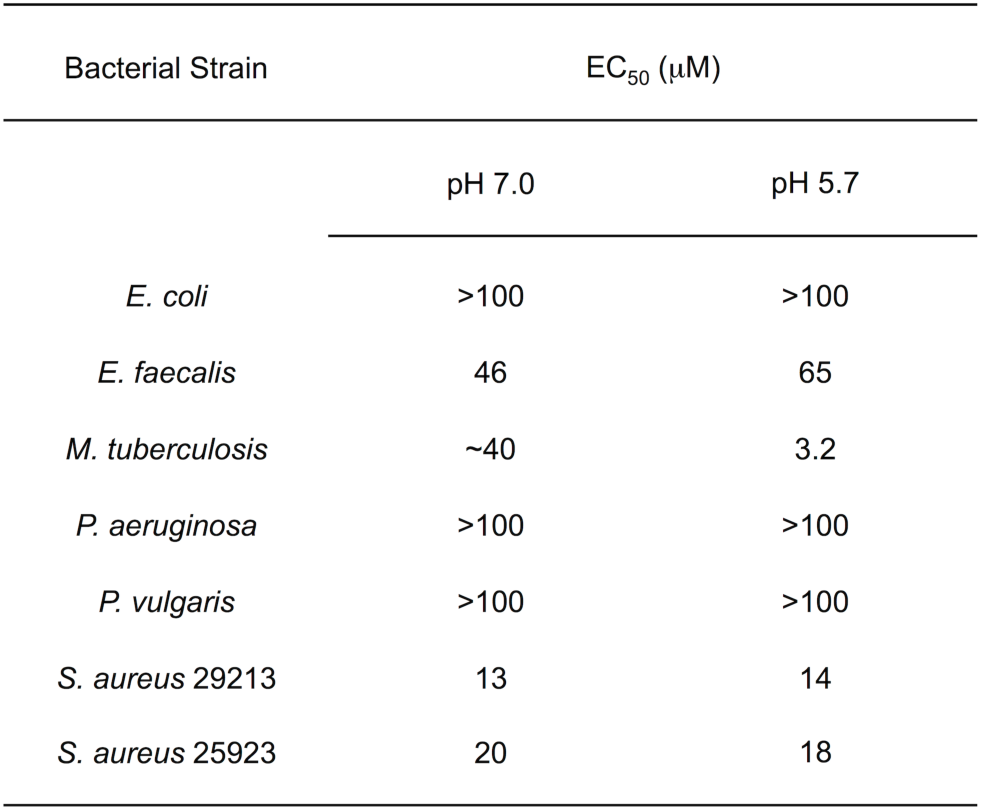
AC2P36 spectrum of activity against Gram-positive and Gram-negative bacteria.

| Bacterial Strain | EC<sub>50</sub> (μM) |  |
| --- | --- | --- |
|  | pH 7.0 | pH 5.7 |
| *E. coli* | >100 | >100 |
| *E. faecalis* | 46 | 65 |
| *M. tuberculosis* | ~40 | 3.2 |
| *P. aeruginosa* | >100 | >100 |
| *P. vulgaris* | >100 | >100 |
| *S. aureus* 29213 | 13 | 14 |
| *S. aureus* 25923 | 20 | 18 |

**Supplemental Table 2.** Transcriptional profiling of Mtb treated with DMSO vs. AC2P36 at pH 7.0.

**Supplemental Table 3.** Transcriptional profiling of Mtb treated with DMSO vs. AC2P36 at pH 5.7.

**Supplemental Table 4.** Transcriptional profiling of DMSO treated Mtb at pH 5.7 vs. pH 7.0.

